# Heterozygote Advantage of a Single-Copy *SNAP18* Truncation Allele Enables Dominant SCN Resistance and Yield Preservation in Soybean

**DOI:** 10.64898/2026.02.24.707632

**Authors:** Dongmei Wang, Shaojie Han, Luying Chen, Liang Wang, Lin Weng, Hui Yu, Chunjie Li, Minghui Huang, Suxin Yang, Xianzhong Feng

## Abstract

Soybean cyst nematode (SCN, *Heterodera glycines*) is the most devastating pathogen threatening global soybean production. The long-term over-reliance on *rhg1*-a/*b* resistance has led to the rapid emergence of virulent SCN populations, creating an urgent demand for novel genetic resources to safeguard soybean yield. Here, we identified a rare gain-of-function allele, *SNAP18_lmm3_*, via map-based cloning of the lesion-mimic mutant (*lmm3*) in the SCN-susceptible Williams 82 (*rhg1-c*) background. This allele encodes a 24-amino-acid C-terminal truncation of SNAP18 and exhibits an unusual mixed dominance genetic profile: the autoimmune-related leaf lesion phenotype is recessive, whereas SCN resistance is dominant. Homozygous *lmm3* plants suffer significant yield losses due to spontaneous cell death, but heterozygous (*LMM3^+/-^*) plants maintain normal growth, agronomic performance, and seed yield comparable to wild-type Williams 82 in SCN-free field environments. Critically, under artificial SCN inoculation in both field trials and controlled environments, *LMM3^+/-^* heterozygotes confer robust resistance, achieving a ∼4-fold yield advantage over SCN-infested Williams 82 and a ∼44% reduction in cyst formation relative to Williams 82. This resistance profile aligns with the broad-spectrum activity of the *lmm3* homozygous mutant, supported by the conserved resistance mechanism of *SNAP18_lmm3_*. Transgenic expression of *SNAP18_lmm3_* driven by the nematode-responsive *HIP* promoter (*HIP_pro_*) potently arrested nematode development at the J2 stage in transgenic hairy roots, as evidenced by the drastically reduced proportion of advanced-stage nematodes. Our findings establish *SNAP18_lmm3_* as a potent, “plug-and-play” genetic resource that circumvents the limitations of traditional dosage-dependent resistance loci. This allele enables the development of high-yielding, SCN-resistant soybean cultivars through marker-assisted selection for the heterozygous state or precision genome editing, providing a practical solution for sustainable SCN management.

## Introduction

Soybean cyst nematode (SCN, *Heterodera glycines* Ichinohe) is a globally devastating plant-parasitic nematode that poses an existential threat to soybean production systems worldwide (Bent 2022). Responsible for 20∼30% of global soybean yield losses, SCN inflicts annual economic damages exceeding US$1.5 billion in the United States alone, with similar impacts in major producing regions such as China and Brazil (Bandara et al. 2020; Bent 2022; Jones et al. 2013). As a sedentary endoparasite, SCN establishes specialized multinucleated syncytia in host root vascular tissues—essential feeding structures that divert host nutrients to support nematode development, thereby crippling soybean growth and productivity (Bent 2022; Yusuf and Bello 2025).

Since the 1990s, genetic improvement has relied heavily on quantitative trait loci (QTL) associated with SCN resistance, with the *Rhg1* (Resistance to *Heterodera glycines 1*) locus emerging as the most widely deployed resource in breeding programs (Concibido et al. 2004). Commercial resistant varieties, such as JTN-5203 (PI 88788-type, *rhg1-b*), IAR1902 SCN (Peking-type, *Rhg1* + *Rhg4*), and N7003CN (*Rhg1* + *Rhg4* + *Rhg5*), have been developed to mitigate SCN damage (Arelli et al. 2015; Carter et al. 2011; Cianzio et al. 2019). The *Rhg1* locus resides on a 31.2-kb tandemly repeated genomic block of chromosome 18, encompassing four genes: *SNAP18* (α-Soluble NSF Attachment Protein 18, alternatively named α-SNAP*_Rhg1_*), *Rhg1-AAT* (putative amino-acid transporter), *WI12_Rhg1_* (wound-inducible 12 domain protein), and *Glyma.18G022600* (function unknown) (Cook et al. 2012, 2014). Among these, *SNAP18*, *Rhg1-AAT*, and *WI12_Rhg1_* are established as major contributors to resistance, while the role of *Glyma.18G022600* remains uncharacterized.

A defining feature of *Rhg1*-mediated resistance is its dosage dependence: increased copy number of the *Rhg1* tandem repeat correlates with elevated transcript levels of resistance-conferring genes in resistant accessions (Cook et al. 2014). Two well-characterized *Rhg1* haplotypes dominate current breeding: (1) *rhg1-a* (low-copy, ≤3 repeats), derived from Peking, which requires epistatic interaction with *Rhg4* (encoding a serine hydroxymethyltransferase with two amino-acid polymorphisms) to confer full resistance against HG Types 0, 1.2.5.7, and 2.5.7 (Patil et al. 2019); and (2) *rhg1-b* (high-copy, ≥ 4 repeats), derived from PI 88788, which provides broad-spectrum resistance independently of *Rhg4*—primarily targeting HG Types 2 and 5, with partial efficacy against HG Type 1.2.5.7 (Patil et al. 2019). In contrast, single-copy *Rhg1* varieties (designated *rhg1-c*), which carry only the canonical *SNAP18* allele, remain highly susceptible to SCN (Cook et al. 2014).

The over-reliance on PI 88788-derived *rhg1-b* alleles in commercial cultivars has severely limited the deployment of alternative resistance strategies, creating a monoculture of resistance that accelerates the evolution of virulent SCN populations (McCarville et al. 2023; Niblack et al. 2008; Rincker et al. 2017; Tylka and Mullaney 2016). Compounding this challenge, SCN’s sexual and promiscuous reproductive strategies enhance genetic diversity, leading to the emergence of new virulent races in major production regions—including the United States and China—that break down traditional *Rhg1*-based resistance (Bent 2022; Chen et al. 2021; Meinhardt et al. 2021; Niblack et al. 2008; Peng et al. 2021). This resistance erosion, coupled with the long-standing bottleneck of yield trade-offs in resistant cultivars, creates an urgent need for novel, durable genetic resources. Early Peking-type (*rhg1-a*) lines exhibited 5.1–14.1% yield losses under SCN-free conditions (Donald et al. 2006), and modern resistant cultivars still suffer modest reductions (2.9–3.9%) compared to susceptible varieties (Rincker et al. 2017).

The reference soybean cultivar Williams 82 (Wm82.a2 genome) encodes five canonical α-SNAP genes (on chromosomes 2, 9, 11, 14, and 18), with the single-copy *SNAP18* on chromosome 18 representing the susceptible *rhg1-c* haplotype (Schmutz et al. 2010). Resistant *rhg1-a* varieties replace this canonical *SNAP18* with ≤ 3 copies of the atypical *SNAP18_LC_* isoform, while *rhg1-b* varieties harbor ≥4 copies of *SNAP18_HC_* alongside one canonical *SNAP18* (Cook et al. 2014; Lee et al. 2015). Additionally, loss-of-function variants of *SNAP11* (chromosome 11) and *SNAP02* (chromosome 2) contribute modestly to resistance in Peking-type soybeans (Lakhssassi et al. 2017; Matsye et al. 2012; Suzuki et al. 2020; Usovsky et al. 2023). However, no single-copy gain-of-function alleles conferring robust, yield-neutral SCN resistance have been identified in *rhg1-c* backgrounds to date.

In this study, we characterize the lesion-mimic mutant *lmm3*—isolated from an EMS-mutagenized Williams 82 population—which harbors a rare gain-of-function allele, *SNAP18_lmm3_*. This allele encodes a 24-amino-acid C-terminal truncation of SNAP18 and exhibits a unique genetic profile of **mixed dominance**: the autoimmune-related leaf lesion phenotype is recessive, while SCN resistance is dominant. Through multi-year, multi-location field trials and controlled-environment assays, we demonstrate that heterozygous (*LMM3*^+/-^) plants provide robust SCN resistance; moreover, this resistance profile aligns with the broad-spectrum activity against HG Types 0, 1.2.5.7, and 2.5.7 observed in the *lmm3* homozygous mutant, all without incurring the agronomic penalties observed in homozygotes Furthermore, we validate the biotechnological utility of *SNAP18_lmm3_* via targeted expression driven by a nematode-responsive *HIP* promoter, which confers high-level resistance in elite soybean backgrounds. Our findings establish *SNAP18_lmm3_* as a potent, single-copy genetic resource that circumvents the biological and breeding limitations of current dosage-dependent resistance sources, offering a “plug-and-play” strategy for sustainable SCN management.

## Results

### Isolation and Phenotypic Characterization of the Soybean *lmm3* Mutant

To identify genes regulating plant immunity and nematode resistance, we screened a soybean mutant library derived from EMS-mutagenesis of the cultivar Williams 82. Among the mutants identified, *lesion-mimic mutant 3* (*lmm3*) develops rust-brown spots on young leaves that expand as the plant grows, leading to necrosis and stem shedding. This phenotype persists throughout its lifecycle (Figure 1A,B). *lmm3* plants are shorter and exhibit reduced 100-seed weight and seed-weight per plant compared to wild-type Williams 82, while showing no statistically significant differences in the number of branches and nodes on the main stem (Figure 1C).

**FIGURE 1.**
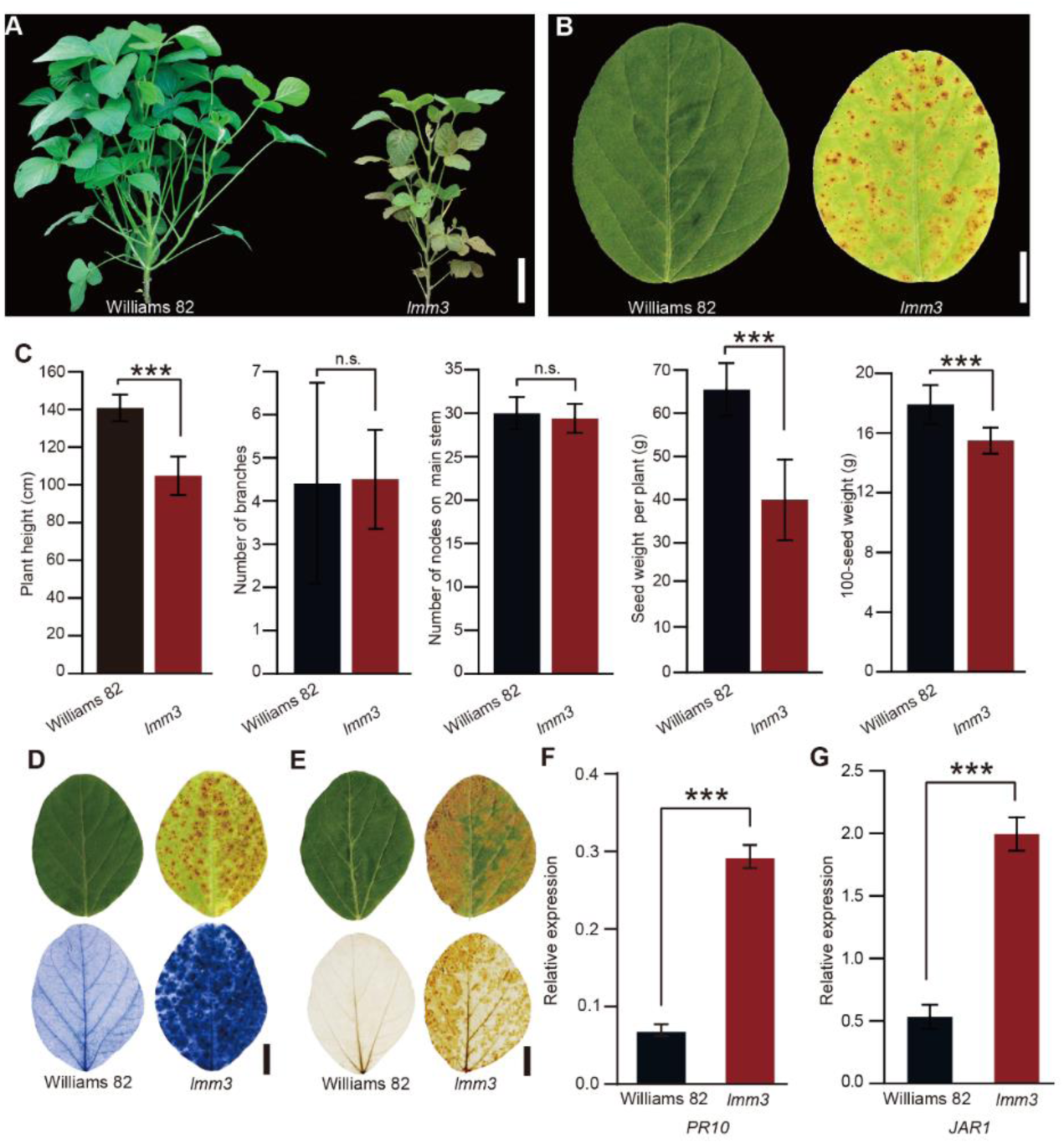
*lmm3* exhibits an autoimmune phenotype in soybean. (A) Representative shoot phenotype of field-grown wild-type c.v. Williams 82 (left) and the *lmm3* mutant (right) at 62 days after germination (DAG) in Changchun, 2023. Scale bar = 10 cm. (B) Representative leaves of Williams 82 (left) and *lmm3* (right) at 28 DAG in a climate chamber in 2023. Scale bar = 1 cm. (C) Yield metrics for field-grown Williams 82 and homozygous *lmm3* mutants. Left to right: Plant height, number of branches, number of nodes on main stem, seed weight per plant (data collected as observed weights following air-drying to constant weight under uniform conditions, with no additional moisture correction) and 100-seed weight for plants grown in Changchun in 2024. The field experiment adopted a randomized complete block design with three blocks and equal replication per genotype. Data represent the mean ±SD (n = 30 for Williams 82, n = 20 for *lmm3*) and asterisks indicate statistically significant differences between means (*** *P* < 0.001, Student’s *t*-test), n.s. denotes non-significant difference. (D) Trypan blue staining on 28-d-old Williams 82 (left) and *lmm3* mutant (right) leaves as a readout for spontaneous cell death. Scale bar = 1 cm. (E) DAB staining on 28-d-old Williams 82 (left) and *lmm3* mutant (right) leaves as a readout for ROS accumulation. Scale bar = 1 cm. (F and G) Expression of *PR10* (F, *Glyma.09G040500*, an SA-responsive marker gene) and *JAR1* (G, *Glyma.19G254000*, a JA-signal-transduction-pathway marker gene) in leaves of 22-d old Williams 82 and *lmm3* mutant plants. Relative expression was normalized to *ACT11*. Data are the mean ± SD of three biological replicates. Asterisks indicate statistically significant differences between means (*** *P* < 0.001, Student’s *t*-test).

Toward understanding the leaf-lesion phenotype, we performed trypan blue staining on the first trifoliolate leaves at the V4 stage four weeks after planting to survey the extent of cell death. This revealed distinct dark-blue spots in suspected necrotic areas of *lmm3*, in contrast to the healthy cells of Williams 82 controls (Figure 1D), suggesting that the lesion-mimic phenotype of *lmm3* is linked to autoimmune-related cell death. Plant immune responses are often associated with accumulation of reactive oxygen species (ROS). DAB staining to assess ROS levels indicated that *lmm3* exhibits enhanced levels of ROS compared to Williams 82 (Figure 1E). Furthermore, *lmm3* displayed an induction of classic defense-hormone-marker genes relative to the wild type, including the SA-responsive *PR10* and JA-responsive *JAR1* (Figure 1F,G). Additionally, mycelial blocks of the major soybean oomycete pathogen *Phytophthora sojae* stain P7076 were inoculated onto the leaves of both the wild type and *lmm3* mutant plants. At 48 hours post-inoculation, lesion areas on the wild-type leaves were statistically significantly larger than those on the *lmm3* leaves, indicating that the mutant has reduced susceptibility to *the pathogen* (Figure S1). These results confirm the autoimmune nature of *lmm3*.

### Map-Based Cloning Identifies *LMM3* as *Glyma.18G022500* Encoding a C-terminally Truncated *α-SNAP*

To identify the causal gene contributing to *lmm3* lesion-mimic phenotype, we first analyzed its inheritance in 550 backcrossed F_2_ plants derived from a cross with Williams 82. The observed 1:3 ratio of mutant to wild-type ratio (122 mutant, 428 wild type) indicated a single, recessive gene (*χ^2^* = 1.06, *df* = 1, *P* = 0.30). Using bulked-segregant analysis, we identified 3,559 SNPs, with a statistically significant peak within the first 5 Mb of chromosome 18 (Figure 2A,B). Within this interval, we identified three genes with a SNP in an annotated exon: a G→A mutation at position 282,116 bp of *Glyma.18G003400* (synonymous mutation), a G→A transition at position 1,027,156 bp of *Glyma.18G014800* (encoding an ATP-dependent RNA helicase, with a glycine-to-threonine change), and a third G→A transition at position 1,645,344 bp of *Glyma.18G022500* (encoding SNAP18). To pinpoint the location of *LMM3*, we crossed *lmm3* with Hedou 12. The F_1_ hybrids exhibited displayed a wild-type leaf appearance without the lesion-mimic phenotype, confirming that the autoimmune-related trait is recessive. F_2_ segregation analysis (193 wild type:56 mutant) matched a 3:1 ratio (*χ^2^* = 0.31, *df* = 1, *P* = 0.58), indicating that the lesion-mimic phenotype is governed by a single recessive mutation. Through map-based cloning using 160 INDEL markers, we localized *LMM3* within a 4.01-Mb segment on chromosome 18, further narrowing it down to a 161-kb interval between markers MOL4708 and MOL4710 (Figure 2C).

**FIGURE 2.**
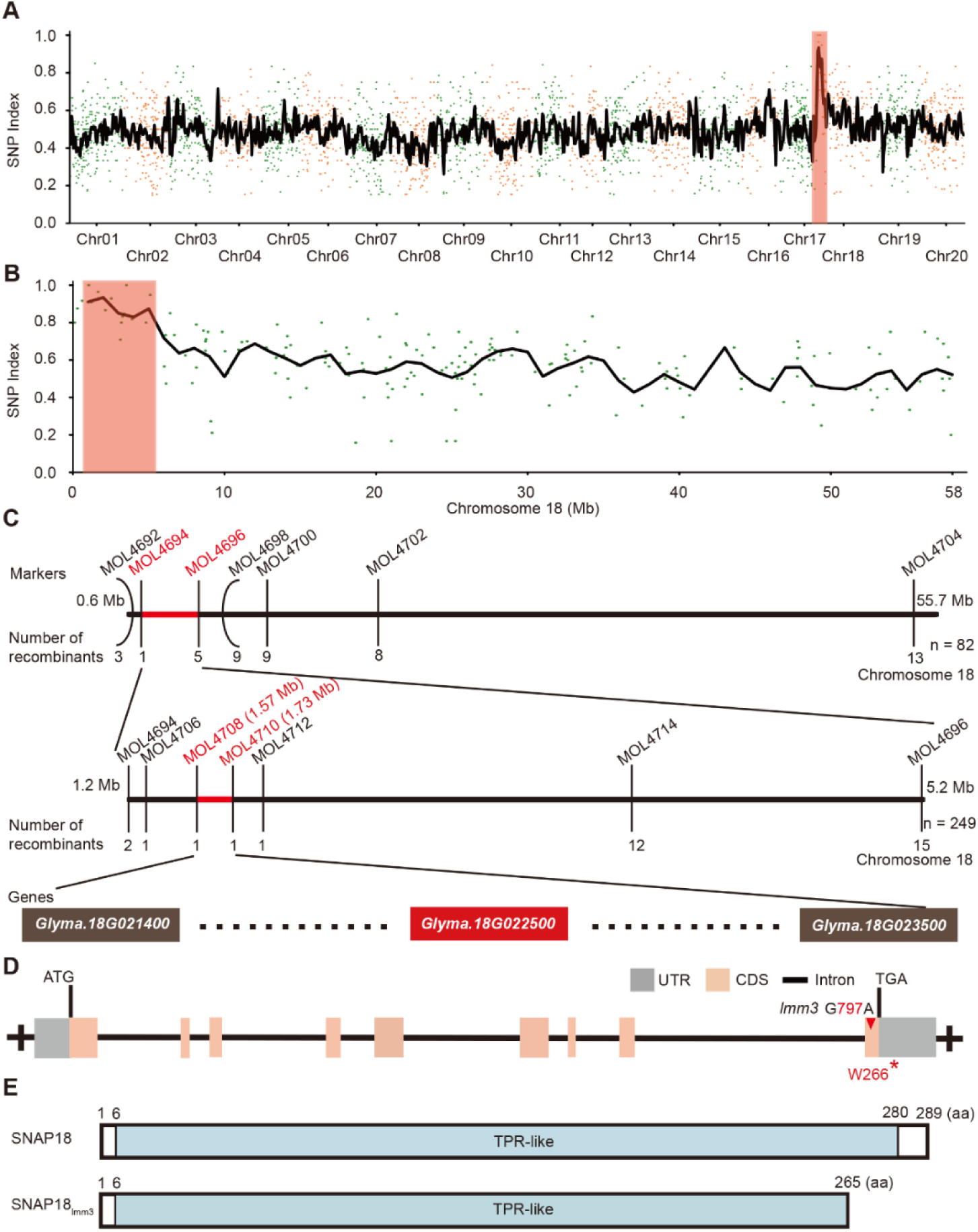
Identification and characterization of *LMM3*. (A and B) Bulked-segregant analysis mapping of *lmm3*. SNP indices across all chromosomes (A) and chromosome 18 (B) of an *lmm3* mutant pool from the BC_1_F_2_ population are shown. The candidate region for *lmm3* is within the first 5 Mb of chromosome 18 (shaded in light red). (C) Map-based cloning of the candidate *lmm3* locus. Top: Coarse positioning of *LMM3* to a 4.01-Mb region flanked by markers MOL4694 and MOL4696. Bottom: Fine-mapping of *LMM3* to a 161-kb region spanned by markers MOL4708 and MOL4710. The number of recombinants detected between the molecular markers is indicated and the number of the plants for map-based cloning is shown adjacent to chromosome 18 in the panel. Not all genes found within the candidate interval are depicted (dotted lines). The candidate gene is depicted within a red box. (D) Genomic structure of *Glyma.18G022500*. The red inverted triangle indicates the mutation site in the *lmm3* mutant. (E) Schematic representations of wild-type Glyma.18G022500 (SNAP18) and the lmm3 mutant protein (SNAP18_lmm3_). SNAP18_lmm3_ is truncated by 24 amino acids due to the introduction of a stop codon resulting from a base substitution in exon 9.

This region included *Glyma.18G022500* and excluded *Glyma.18G003400* and *Glyma.18G014800*. The G→A transition at position 1,645,344 bp of *Glyma.18G022500* (*SNAP18*) causes a mutation at nucleotide-position 797 of the coding sequence, resulting in a premature stop codon introduced at amino-acid 266 in exon 9 (Figure 2D,E). By combining bulked-segregant analysis and map-based cloning, we identified *Glyma.18G022500* as the *LMM3* gene. *Glyma.18G022500* is linked to SCN resistance (Bayless et al. 2016; Cook et al. 2012; Usovsky et al. 2023), and its exact role in the context of the *lmm3* phenotype warrants further investigation.

Consistent with monogenic recessive inheritance, genotyping of 550 BC_1_F_2_ and 249 Hedou 12-derived F_2_ individuals showed strict co-segregation between the *Glyma.18G022500* G→A mutation (*lmm3*) and the lesion-mimic phenotype, with all 122 BC_1_F_2_ mutant-phenotype plants homozygous for the *lmm3* mutation and all 428 wild-type-phenotype plants carrying at least one wild-type *LMM3* allele. Similarly, 56 Hedou 12-derived F_2_ mutant plants were homozygous for *lmm3*, and 193 wild-type plants were *LMM3*/– (χ² = 1.06 and 0.31, respectively, *P* > 0.3). No recombinant individuals were detected within the fine-mapped 161-kb interval, confirming tight linkage between the mutation and phenotype. These results confirm *lmm3* as the causal locus, minimizing the possibility of background effects confounding the phenotype.

### Genetic Complementation Confirms *Glyma.18G022500* as the Functional *LMM3* Locus

To verify whether loss of *Glyma.18G022500* leads the *lmm3* mutant phenotype, we aimed to genetically complement the mutant by stably introducing the wild-type *LMM3* coding sequence from Williams 82, fused to green-fluorescent protein (GFP), driven by a 3.12-kb *LMM3* promoter fragment (designated *LMM3_pro_:LMM3-GFP*, Figure 3A,B). Among 22 T_0_ transgenic lines PCR-positive for the *Bar* herbicide-resistance gene (Figure S2A), eight T_1_ progeny restored the wild-type phenotype. Three T_1_ lines, which exhibited no signs of necrosis in their leaves, were further characterized (Figure 3C). PCR and sequencing confirmed these lines carry both the wild-type (from the transgene) and *lmm3* mutant genotypes (Figure S2B-D), and expression of LMM3-GFP was validated by anti-GFP western blotting (Figure S2E). Trypan blue staining indicated no detectable cell death, and there were no signs of ROS over-accumulation or reddish-brown deposits in necrotic areas of *LMM3_pro_:LMM3-GFP* leaves (Figure 3D,E). These findings strongly suggest that *Glyma.18G022500* encodes LMM3. Consequently, these results demonstrate that the 24-amino-acid C-terminal truncation in SNAP18 is the causal mutation underlying the autoimmune-related traits in *lmm3*.

**FIGURE 3.**
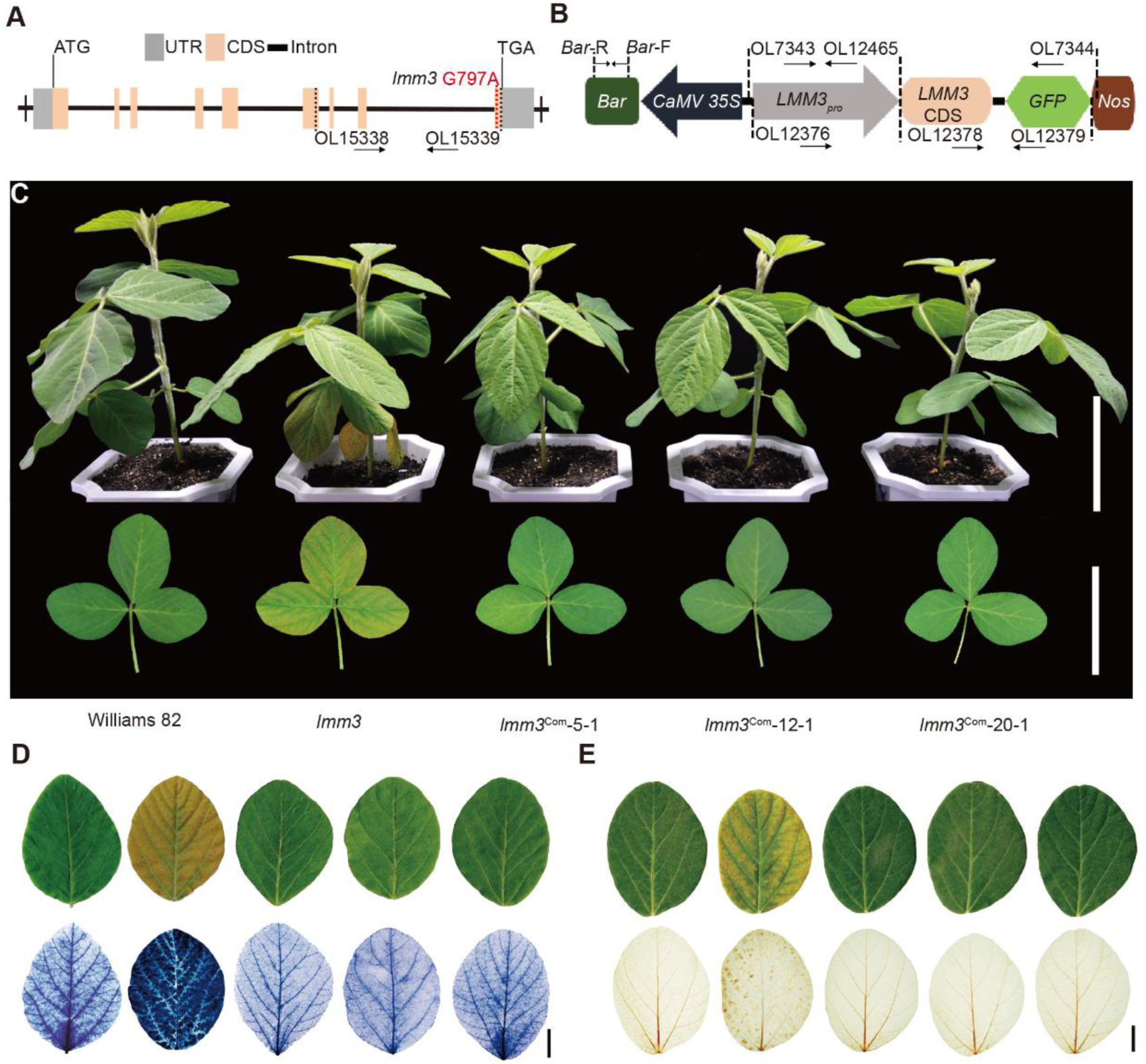
Genetic complementation confirms that *Glyma.18G022500* is the causal gene for the *lmm3* phenotype. (A) Genomic structure of *Glyma.18G022500*. The red dashed line represents the single-nucleotide mutation at nucleotide position 797 in exon 9 in the *lmm3* mutant. The two black dashed lines indicate binding sites for primers OL15338 and OL15339, which amplify the genomic sequence containing three introns in Williams 82 or *lmm3*, or amplify a truncated coding sequence of *Glyma.18G022500* in the plasmid from the complementation construct. The black arrows indicate the direction of the corresponding primers. (B) Summary of the T-DNA region of the *LMM3_pro_:LMM3*–*GFP* complementation construct. Black dashed lines indicate amplicons from the respective primers and the corresponding amplification directions are marked by black arrows. (C) Shoot (top) and first-trifoliolate-leaf (bottom) phenotypes of Williams 82, *lmm3* and three genetically independent T_1_ transgenic complementation events (*lmm3*^Com^-5-1, *lmm3*^Com^-12-1 and *lmm3*^Com-^20-1) harboring *LMM3_pro_:LMM3–GFP* constructs in the *lmm3* mutant background. Plants were photographed after 4 weeks of cultivation in a growth chamber. Scale bars = 10 cm (top) and 5 cm (bottom). (D) Trypan blue staining in leaves as a readout of spontaneous cell death for 30-d-old Williams 82, *lmm3* and the three independent T_1_ transgenic *LMM3_pro_:LMM3*–*GFP* complementation lines. Scale bar = 1 cm. (E) DAB staining to detect ROS accumulation in leaves of 30-d-old Williams 82, *lmm3* and the three independent T_1_ transgenic *LMM3_pro_:LMM3–GFP* complementation lines. Scale bar = 1 cm.

### SNAP18_lmm3_ Confers Broad-Spectrum Resistance Against Multiple SCN Races

*Glyma.18G022500*, also known as *SNAP18*, encodes an α-SNAP that physically interacts with NSF in the SNARE complex to facilitate vesicle trafficking (Hay and Scheller 1997; Huang et al. 2019). α-SNAPs derived from *rhg1* loci, with varying copy numbers, represent naturally occurring *SNAP18* alleles (Figure 4A), and these copy-number-variant alleles contribute to SCN resistance. The *lmm3* mutant encodes an abnormal α-SNAP with a C-terminal truncation of 24 amino acids, designated SNAP18_lmm3_ (Figure 2D,E).

**FIGURE 4.**
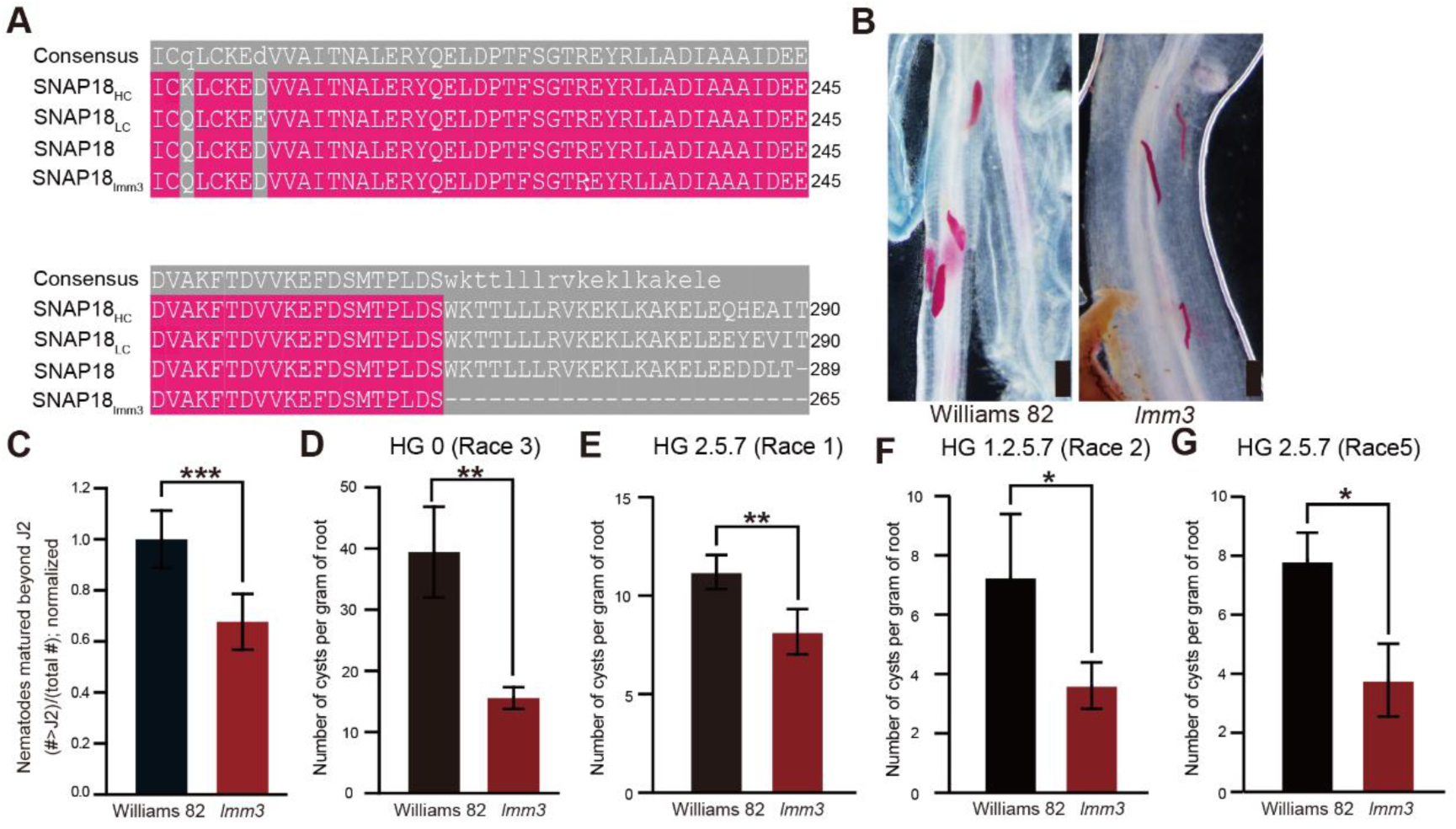
SNAP18_lmm3_ confers soybean broad-spectrum resistance against multiple SCN races. (A) C-terminal amino-acid polymorphisms in SNAP18, SNAP18_HC_, SNAP18_LC_ and SNAP18_lmm3_ isoforms. Williams 82 encodes a canonical SNAP18 protein and is susceptible to SCN. (B) Representative images of fuchsin-stained *H. glycines* observed in roots of Williams 82 and *lmm3* plants at 15 days post-infection (dpi). Scale bar = 200 μm. (C) Analysis of SCN development beyond J2 stage in SCN-susceptible Williams 82 and *lmm3* roots at 15 dpi. Data are the mean ±SD of n = 10 plants per replicate. \*\*\**P* < 0.001, Student’s *t*-test. (D) Quantification of cysts per gram of root in SCN-susceptible Williams 82 and *lmm3* plants at 30 dpi. Data are the mean ± SEM of n = 10 plants per replicate. \*\**P* < 0.01, Student’s *t*-test. (E-G) Quantification of cysts per gram of root in Williams 82 and *lmm3* plants at 45 dpi, infected by different SCN races (Race 1: HG 2.5.7; Race 2: HG 1.2.5.7; Race 5: HG 2.5.7) in a climate chamber. Data are presented as mean ±SD (n = 4 plants). \**P* < 0.05, \*\**P* < 0.01, Student’s *t*-test.

To elucidate the function of SNAP18_lmm3_ in SCN resistance, we inoculated both Williams 82 and *lmm3* plants with nematodes using methods developed by Cook and Liu (Cook et al. 2012; Liu et al. 2012). At 15 days post-infection (dpi), nematodes parasitizing wild-type plants had progressed to the parasitic third stage (par-J3), characterized by a thick and short appearance. In contrast, the nematodes in *lmm3* predominantly remained in the parasitic second stage (par-J2) with a thin and elongated morphology (Figure 4B). The number of nematodes beyond the J2 stage in *lmm3* roots were statistically significantly reduced compared to those in Williams 82 (Figure 4C). By 30 dpi, most nematodes entered the reproductive stage, leading to the formation of cysts in both host genotypes. However, the number of cysts per gram or root on *lmm3* was statistically significantly lower than that on Williams 82 (Figure 4D). At 8 days without SCN, fresh root weight and gross morphology did not differ between Williams 82 and *lmm3* (Figure S3A). Under SCN challenge, Williams 82 showed ∼150% greater root fresh weight than *lmm3* at 7 dpi, whereas by 30 dpi *lmm3* exceeded Williams 82 by ∼54% (Figure S3B). These dynamics are consistent with greater SCN tolerance in *lmm3*, supporting a role of SNAP18_lmm3_ in soybean defense. Results derive from independent biological replicates with n = 10 plants per genotype in each experiment. Taken together, these results indicate that *lmm3* exhibits greater tolerance to SCN challenge, highlighting the role of SNAP18_lmm3_ in soybean defense against this pathogen. Controlled-environment assays further revealed that SNAP18_lmm3_ confers broad-spectrum resistance across diverse SCN populations, including HG Type 0 (Race 3), Race 1 (HG 2.5.7), Race 2 (HG 1.2.5.7), and Race 5 (HG 2.5.7) (Figure 4D-G).

### Heterozygous *LMM3^+/-^* Plants Maintain Growth and Yield in SCN-Free Environments

Williams 82, although not considered a high-yielding cultivar by modern standards, has historically been widely used as an elite background due to its agronomic stability. If provided with effective SCN resistance, it could maintain its favorable yield potential under nematode pressure, comparable to or exceeding that of SCN-resistant varieties derived from PI88788 (*rhg1-b*) or Peking (*rhg1-a*) (Bent 2022). Importantly, reducing SCN susceptibility in Williams 82 through a single gene without yield trade-offs remains a desirable outcome.

*SNAP18_lmm3_* potentially offers a distinct benefit in resistance breeding as a single-copy, dominant source of SCN suppression. Unlike the homozygous *lmm3* mutant, which suffers from severe autoimmune-related growth inhibition, the heterozygous state appears to decouple the beneficial resistance from the detrimental pleiotropic effects. To explore breeding utility, we profiled field-grown wild-type Williams 82 and both heterozygous and homozygous *lmm3* plants. Plants were genotyped using CAPS assay (Figure S4). Yield was defined as single-plant seed weight at maturity. Across two field environments arranged as randomized complete block designs with three blocks, *LMM3*^+/-^ heterozygotes showed seed yield comparable to Williams 82, whereas homozygous *lmm3* exhibited a reduction. Plot-level yields standardized to 13% moisture are provided in Table S1, and no significant genotype-by-environment interaction was detected for Williams 82 versus *LMM3*^+/-^ heterozygotes under the tested environments (Figure 5A,B and Table S1). Seed-weight per plant, 100-seed weight, and plant height for heterozygotes were statistically non-significantly different from Williams 82 (Figure 5C-E).

**FIGURE 5.**
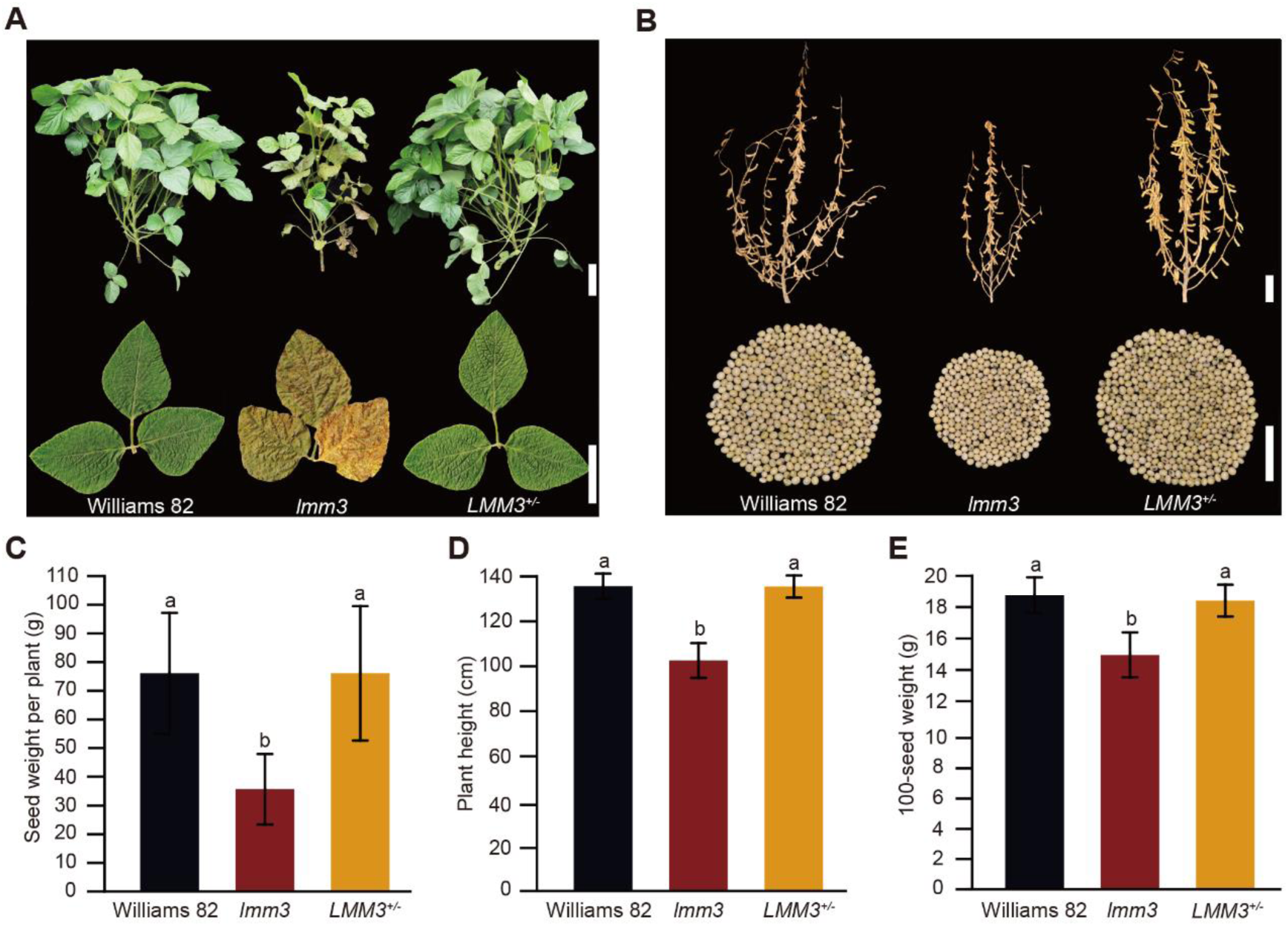
Heterozygous *LMM3^+/-^* plants maintain growth and yield in SCN-free environments. (A) Representative phenotypes of field-grown Williams 82, *lmm3* mutant and *LMM3^+/-^*heterozygous plants observed 72 days post-planting at the R2 stage. Scale bars = 10 cm for shoots and 5 cm for trifoliolate leaves. (B) Phenotypes of mature field-grown Williams 82, *lmm3* and heterozygous *LMM3^+/-^* plants with shoots with leaves removed (top) and seeds (bottom). Scale bars for shoots = 10 cm and 5 cm for seed pools. (C*–*E) Yield metrics for field-grown mature Williams 82, *lmm3* and heterozygous *LMM3^+/-^*plants. Seed-weight per plant (C), plant height (D) and 100-seed weight (E). Data represent the mean ±SD of n = 30 plants. One-way ANOVA, different letters represent *P* < 0.05. Plants in panels (A*–*E) were field-grown in Changchun in 2023 (43.88°N, 125.35°E) in SCN-free environments under a randomized complete block design with three blocks and equal replication per genotype. For seed-weight per plant, data were collected as observed weights following air drying to constant weight under uniform conditions, with no additional moisture correction.

These results suggest that a single normal copy of *SNAP18* is sufficient to compensate for the cytotoxic effects caused by *SNAP18_lmm3_* (Figure 5A-E). While the homozygous mutant exhibits systemic cell death, the heterozygous plants maintain a healthy phenotype and normal agronomic performance, facilitating the maintenance of high yield potential in SCN-free environments. The detailed cellular mechanisms allowing for this balanced homeostasis are explored in our companion study.

### *LMM3^+/-^* Heterozygotes Confer Robust SCN Resistance while Protecting Seed Yield

To evaluate the resistance efficacy of the *SNAP18_lmm3_* allele in a heterozygous state, we first monitored SNAP18 abundance under both uninfected and SCN-infected conditions. Upon SCN challenge, *LMM3^+/-^* heterozygous roots showed a significant induction of SNAP18 protein, a pattern similar to that observed in homozygous *lmm3* mutants (Figure 6A).

**FIGURE 6.**
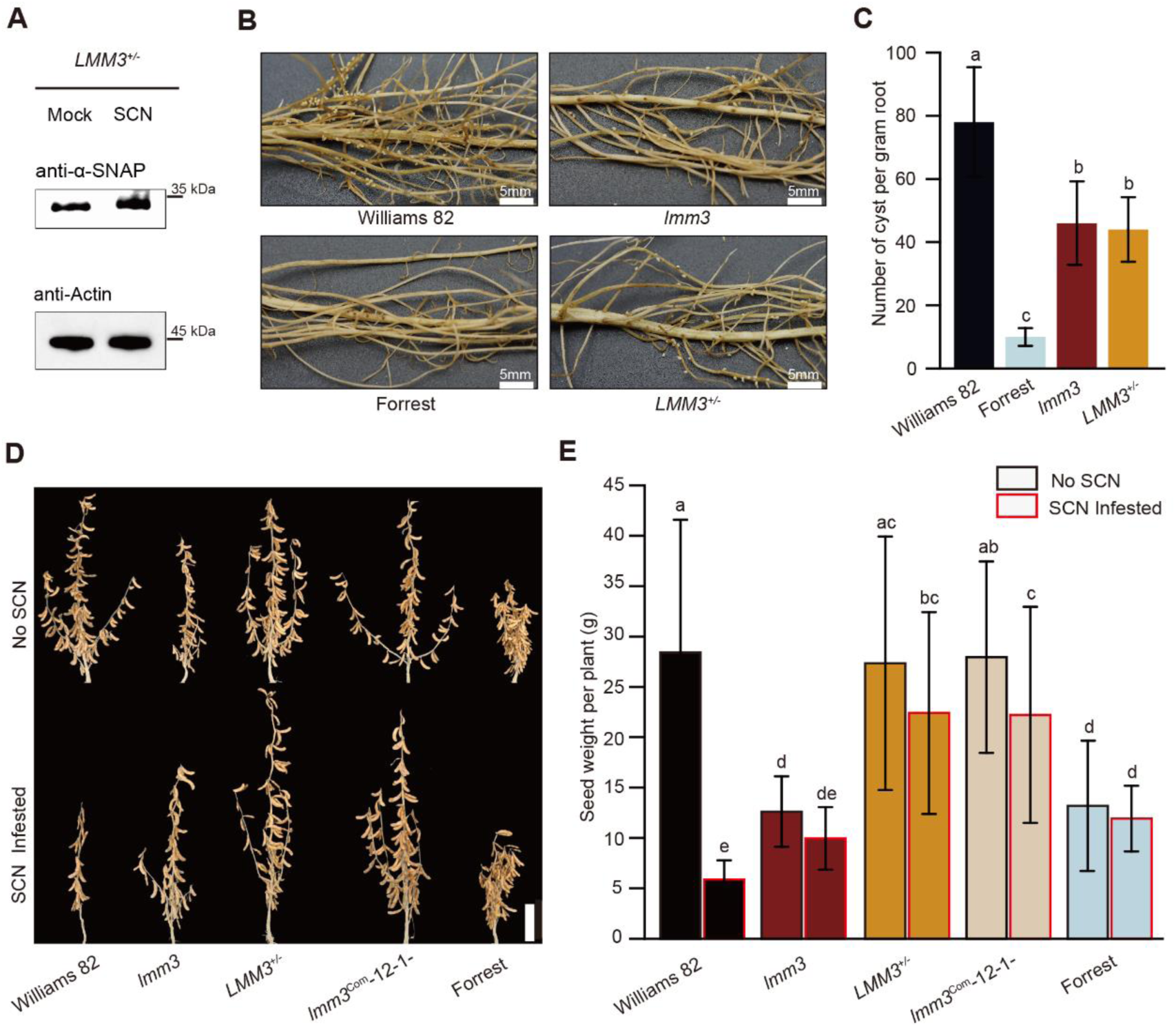
Genetic and functional characterization of SNAP18_lmm3_-mediated resistance to SCN in soybean. (A) Western blot of α-SNAP abundance in root samples from uninfected and SCN-infected heterozygous *LMM3^+/-^* plants harvested 3 dpi, with actin used as the loading control. (B) Representative images of infected roots from Williams 82, Forrest, *lmm3*, and heterozygous *LMM3^+/-^* plants grown in a climate chamber under SCN challenge at 45 dpi. Scale bars = 5 mm. (C) Quantification of cysts at 45 dpi on roots of Williams 82, Forrest, *lmm3* and heterozygous *LMM3^+/-^*plants grown in a climate chamber under SCN challenge. Data are presented as mean ± SD (n ≥ 5 plants). One-way ANOVA, different letters represent *P* < 0.05. (D) Representative phenotypes of mature field-grown Williams 82, *lmm3* mutant, *LMM3^+/-^* heterozygous plants, T_2_ transgenic complementation plants (*lmm3*^Com^-12-1-, harboring *LMM3_pro_:LMM3–GFP* constructs in the *lmm3* mutant background) and Forrest, with or without SCN infestation. Scale bar = 10 cm. (E) Seed-weight per plant corresponding to the plants shown in panel (D). Data represent the mean ±SD of n = 45 plants. Two-way ANOVA, different letters represent *P* < 0.05. Plants in panels (D) and (E) were field-grown in Tonglu in 2024 (29.835°N, 119.603°E) under a randomized complete block design with three blocks and equal replication per genotype under both SCN-inoculated and non-inoculated conditions. For seed-weight per plant, data were collected as observed weights following air drying to constant weight under uniform conditions, with no additional moisture correction.

To quantify this resistance, plants propagated in the growth chamber were inoculated with J2-stage nematodes, and 45 d later, prior to cyst counting, representative images of infected roots showed numerous cysts on the roots of the susceptible SCN variety Williams 82. In contrast, the *lmm3* mutant and heterozygous (*LMM3*^+/-^) roots exhibited statistically significantly fewer cysts compared to Williams 82. Only a few cysts were visible on the roots of the SCN-resistant control variety Forrest (Figure 6B). Heterozygous plants exhibited comparable reductions in SCN susceptibility to homozygous *lmm3* mutants under controlled conditions (Figure 6C). Even after approximately two generations of SCN growth, heterozygosity still provided similar resistance to that of the homozygous *lmm3* mutant carrying two *SNAP18_lmm3_* alleles.

To clarify both per-plant and plot-level yield performance, as well as potential environmental effects, we conducted two field trials arranged as randomized complete block designs with three blocks: Changchun 2023 without SCN inoculation and Tonglu 2024 with paired inoculated and non-inoculated plots (Figure 6D,E; Figure S5 and Table S1). Under SCN-free conditions, heterozygous plants produced yields comparable to Williams 82, indicating no negative impact on yield potential. In SCN-infected conditions, heterozygous plants demonstrated reduced susceptibility to SCN, maintaining higher seed weights with only a slight decline compared to their performance in SCN-free environments. In contrast, Williams 82 exhibited a significant yield reduction when challenged with SCN. Notably, similar trends were observed in T_2_ transgenic complementation plants (*lmm3*^Com^-12-1-), which phenotypically resemble heterozygotes with stable inheritance (Figure 6D,E; Figure S5 and Table S1). These findings suggest that heterozygous plants carrying *SNAP18_lmm3_* provide effective yield protection and reduced susceptibility to SCN, underscoring their strong potential for breeding programs targeting SCN resistance. Notably, *SNAP18_lmm3_*-mediated resistance differs from classical SCN strategies in both genetic architecture and mechanism. Multi-copy *Rhg1* relies on increased a-SNAP dosage together with additional locus components, and Peking-type resistance requires epistasis between *Rhg1* and *Rhg4*. By contrast, *SNAP18_lmm3_* is a single-copy truncation allele that shows rapid local accumulation at SCN feeding sites. This profile is consistent with the observation that heterozygotes retain resistance without lesion-mimic symptoms or detectable yield penalties in the Williams 82 and Forrest backgrounds under the conditions tested.

### Biotechnological Deployment of *SNAP18_lmm3_* via SCN-Responsive Targeted Expression

To overcome hybrid-segregation challenges, we used genetic engineering to introduce *SNAP18_lmm3_* into transgenic soybeans and assess SCN resistance. We used hairy-root transformation for its speed and phenotypic insight. The SCN-responsive *HIP* promoter (Kandoth et al. 2011) drove the expression of *SNAP18_lmm3_* (Figure 7A). After validating the promoter’s response to SCN challenge (Figure 7B), we infected transgenic roots with nematodes (Figure 7C). At 12 dpi, nematodes typically progressed to the thick, short parasitic third stage (par-J3) on control roots, whereas those found in the *HIP_pro_:lmm3* transgenic roots, regardless of the Williams 82 or Forrest background, predominantly bore thinner, elongated, parasitic second-stage (par-J2) nematodes (Figure 7D). This resulted in a markedly lower advancement of SCN beyond the J2 stage in *HIP_pro_:lmm3* roots, highlighting enhanced nematode resistance across different elite backgrounds (Figure 7E). This confirms the viability of introducing *SNAP18_lmm3_* into cultivars through genetic engineering to promote resistance.

**FIGURE 7.**
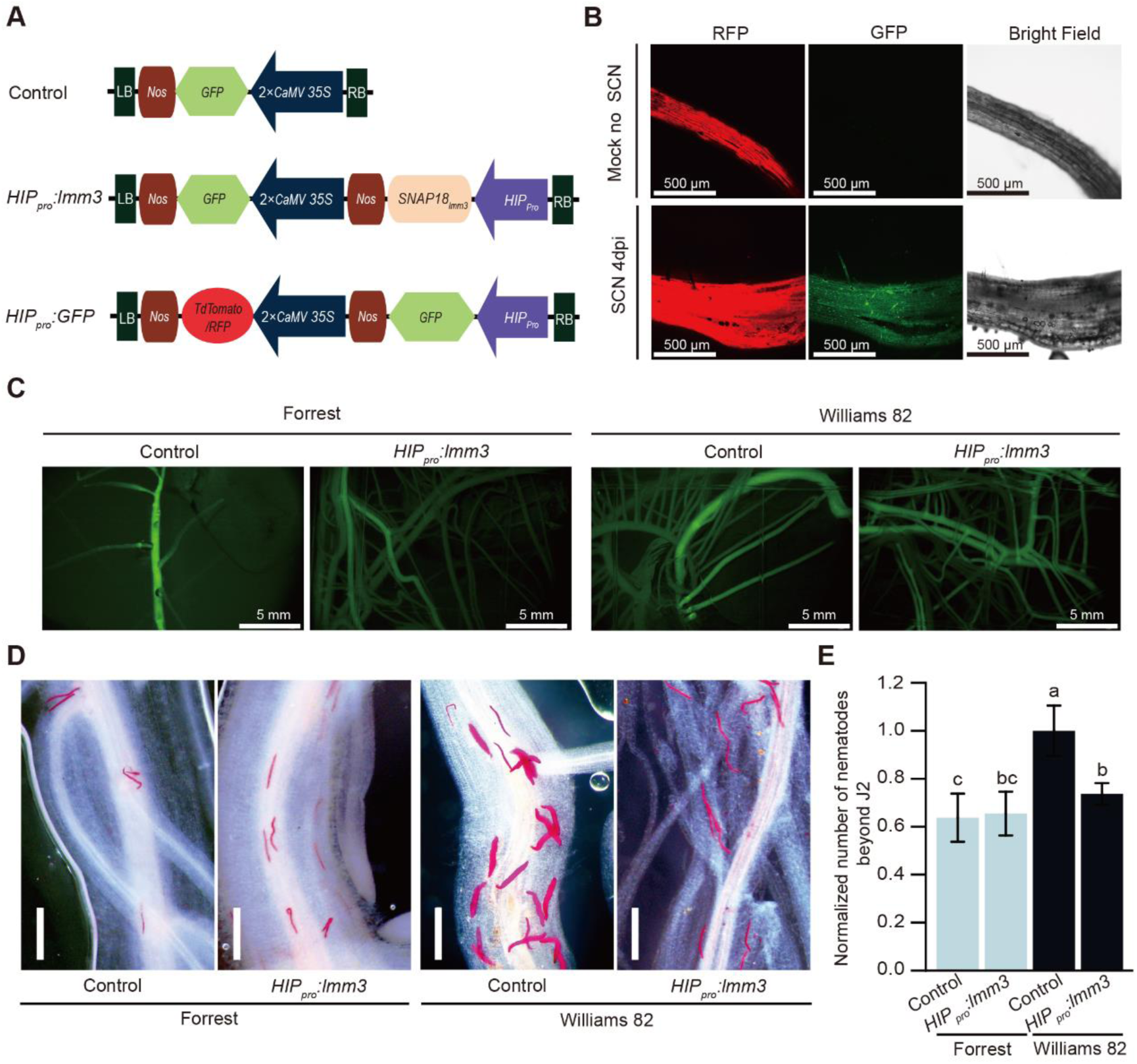
Transgenic expression of *SNAP18_lmm3_* in soybean hairy roots confers SCN suppression. (A) Summary of T-DNA regions of constructs used for transformation. A control plasmid was designed based on the pCAMBIA3301T backbone, with the GFP selectable marker driven by the *35S* promoter. *HIP_pro_:lmm3* plasmid: A 1-kb *HARPIN-INDUCED PROTEIN* (*HIP, Glyma.03G201100*) promoter region was used to control the expression of *SNAP18_lmm3_*. For facile selection of transgenic hairy roots, an independent *GFP* cassette is also driven by the 2 ×*CaMV 35S* promoter. *HIP_pro_:GFP* plasmid: This employs the same *HIP* promoter to regulate *GFP* expression, providing a means to gauge SCN responsiveness. (B) Epifluorescence microscopy of transgenic hairy roots bearing the *HIP_pro_:GFP* construct either exposed to SCN or mock treatment. GFP fluorescence is detectable by 4 dpi. Scale bar = 500 μm. (C) Epifluorescence microscopy with a 495-nm band-pass filter of transgenic hairy roots differentiated from native roots, of a representative GFP-positive transformed root. Scale bar = 5 mm. (D) Representative fuchsin-stained *H. glycines* on 12-dpi transgenic hairy roots featuring control or *HIP_pro_:lmm3* constructs in the c.v. Forrest (SCN resistant) and c.v. Williams 82 (SCN susceptible) wild-type backgrounds. *GFP* driven by the *35S* promoter was used as a transformation control. Scale bar = 500 μm. (E) SCN demographics in transgenic hairy roots expressing the transformation control or *HIP_pro_:lmm3* constructs in c.v. Forrest and c.v. Williams 82 backgrounds as in (D). Data are the mean ±SD of n = 9 hairy roots. Statistical significance determined by two-way ANOVA, with differing letters indicating *P* < 0.05.

## Discussion

### The Unique Mixed-Dominance Profile of *SNAP18_lmm3_* Revolutionizes Resistance Breeding

The discovery of the *lmm3* lesion-mimic mutant marks a significant advancement in SCN resistance research in soybean. This mutant harbors a 24-amino-acid C-terminal truncation of SNAP18, designated SNAP18_lmm3_. This allele’s most distinctive feature is its **mixed dominance** genetic pattern: the deleterious autoimmune leaf lesion phenotype is recessive, while SCN resistance is dominant. Such decoupling of beneficial and detrimental traits is rare in plant genetics, with few analogs (e.g., the *Curly* mutation in *Drosophila*) (Lindsley and Grell 1972), and offers unprecedented breeding advantages. Our multi-location field trials confirm that homozygous *lmm3* plants suffer stunted growth and yield losses due to systemic cell death, but heterozygous (*LMM3^+/-^*) individuals retain the elite agronomic performance of Williams 82 in SCN-free environments—with seed yield (2399 kg/ha), 100-seed weight, and plant height statistically indistinguishable from the wild type. Under artificial SCN inoculation in both field trials and controlled environments, these heterozygotes outperform susceptible cultivars by nearly four-fold (1927 vs. 517 kg/ha) (Table S1) and reduce cyst formation by 44% (Figure 6C), resolving the long-standing yield-resistance trade-off that has plagued soybean breeding for decades (Donald et al. 2006; Rincker et al. 2017).

This mixed dominance likely arises from the dosage-dependent balance between SNAP18_lmm3_’s cytotoxicity and the wild-type SNAP18’s compensatory function. In heterozygotes, a single copy of canonical *SNAP18* suffices to maintain vesicular trafficking homeostasis (as explored in our companion mechanistic study), while SNAP18_lmm3_ accumulates locally at SCN feeding sites to trigger resistance. In homozygotes, the absence of wild-type SNAP18 leads to unrestricted SNAP18_lmm3_ accumulation, overwhelming cellular detoxification mechanisms and inducing systemic necrosis. This genetic architecture simplifies breeding: marker-assisted selection (MAS) for the heterozygous state eliminates the need for complex genotypic-phenotypic validation, enabling rapid introgression into elite germplasm.

### *SNAP18_lmm3_*: A Single-Locus Alternative to Complex *Rhg1*-Mediated Resistance

Traditional SCN resistance strategies rely on either gene copy expansion (*rhg1-b*) or epistatic crosstalk between multiple loci (*rhg1-a* + *Rhg4*) (Bent 2022; Cook et al. 2014). These approaches suffer from inherent limitations: *rhg1-b*’s dosage dependence increases breeding complexity and risks pleiotropic effects, while *rhg1-a*’s requirement for *Rhg4* restricts its deployment to genetic backgrounds carrying this interacting locus. In contrast, *SNAP18_lmm3_* is a single-copy gain-of-function allele that confers robust resistance independently of additional loci or copy number variation—even in the susceptible *rhg1-c* background (Williams 82). This simplicity eliminates linkage drag and dosage-dependent regulation, addressing key bottlenecks in current breeding pipelines.

The allele’s efficacy in Williams 82 highlights its ability to target a conserved vulnerability in SCN parasitism: the nematode’s reliance on functional syncytia for nutrient acquisition. As demonstrated in our companion study, SNAP18_lmm3_ disrupts vesicular trafficking by switching binding specificity from NSF to ATG8f, triggering localized autophagic cell death at feeding sites. This mechanism bypasses the need for traditional resistance backgrounds, making *SNAP18_lmm3_* compatible with diverse soybean germplasm. Furthermore, its broad-spectrum activity against HG Types 0, 1.2.5.7, and 2.5.7 (virulent races that break down *rhg1-b* resistance), positions it as a critical resource for managing evolving SCN populations (Chen et al. 2021; Meinhardt et al. 2021).

### Precision Engineering via Nematode-Responsive Promoters Mitigates Pleiotropic Risks

To fully exploit *SNAP18_lmm3_*’s potential while avoiding hybrid segregation challenges, we developed a “plug-and-play” biotechnological strategy using the nematode-responsive *HIP* promoter (*Glyma.03G201100*). This promoter is specifically activated at SCN feeding sites (syncytia) upon infection, driving localized *SNAP18_lmm3_* expression and avoiding the systemic cytotoxicity associated with constitutive promoters (Kandoth et al. 2011).

We evaluated this spatio-temporal control system using transgenic hairy root assays in the elite backgrounds Williams 82 and Forrest. The targeted expression of *SNAP18_lmm3_* conferred robust SCN resistance in these composite plants compared to controls. These results demonstrate that mimicking the syncytia-specific accumulation of the natural mutant is a viable approach for deploying cytotoxic resistance genes. This targeted strategy provides a robust framework for future engineering of SCN-resistant soybean varieties without compromising overall root health or plant development.

This targeted expression approach builds on prior advances in plant biotechnology (e.g., pathogen-responsive promoters for disease resistance; Wei et al. 2024) but is uniquely tailored to SCN’s biology. By restricting SNAP18_lmm3_ accumulation to syncytia, we decouple its defensive function from growth inhibition, providing a blueprint for engineering resistance in other crops without yield trade-offs. Notably, the *HIP* promoter’s conservation across soybean varieties (Kandoth et al. 2011) ensures this strategy’s broad applicability, further enhancing SNAP18_lmm3_’s translational value.

While our findings establish *SNAP18_lmm3_* as a transformative genetic resource, several avenues warrant further investigation to maximize its impact. First, multi-location trials across diverse agroecosystems are needed to validate yield stability and resistance efficacy under field-realistic conditions, with long-term monitoring to assess durability against emerging SCN races, given significant genotype-by-environment interactions (Rincker et al. 2017). Second, stacking *SNAP18_lmm3_* with existing resistance QTLs (e.g., *Rhg4*, *Rhg5*) or transgenic traits (e.g., *Bt*) could enhance broad-spectrum pest management. Its distinct mechanism (vesicular trafficking disruption + autophagic cell death) versus *Rhg1*-mediated resistance (SNARE complex impairment; Bayless et al. 2016) may yield synergistic effects. Third, exploring the molecular basis of its mixed dominance could inform synthetic resistance alleles in other crops, while investigating cross-resistance to pathogens like *Phytophthora sojae* (Figure S1) expands utility beyond SCN. Finally, translating to breeding requires high-throughput genotyping tools (e.g., KASP assays) to complement existing CAPS markers (Figure S4), and precision genome editing (e.g., CRISPR-Cas9) to recreate the 24-amino-acid truncation in elite cultivars, accelerating resistance development. In summary, *SNAP18_lmm3_*’s mixed dominance enables heterozygotes to confer robust, broad-spectrum SCN resistance without yield penalties, operating independently of gene copy expansion or epistatic interactions. Validated via nematode-responsive promoter, it offers a “plug-and-play” solution for sustainable SCN management while advancing understanding of plant immunity and cellular homeostasis.

## Materials and Methods

### Plant Materials and Growth Conditions

The soybean cultivars Williams 82, Hedou 12, and Forrest (*rhg1-a*) were used. The *lmm3* mutant was isolated from an EMS-mutagenized Williams 82 population (Gao et al. 2020; Wang et al. 2020) and stabilized through four generations of selfing and backcrossing. Hedou 12 served as the mapping parent for the F_2_ population. For controlled assays, plants were grown in a climate chamber at 26 °C, 50% humidity, and a 14-hour photoperiod.

### Field Experiments and Yield Determination

Field trials were conducted across two years in two locations: Changchun (43.88°N, 125.35°E) in 2023∼2024 (SCN-free) and Tonglu (29.835°N, 119.603°E) in 2024 (paired SCN-inoculated and non-inoculated plots). All experiments followed a randomized complete block design (RCBD) with three replicates.

Plants were grown with 60 cm between rows and 20 cm between plants (approximately 87,500 plants/ha). For yield metrics, 15 consecutive plants per plot (45 plants per genotype total) were sampled at the R8 stage. Seed weight per plant, 100-seed weight, and plant height were recorded. All seed weights were standardized to a 13% moisture content using a KETT PM8188A meter.

### Nematode-Infection Assays

Nematode-infection assays for soybean plants in the greenhouse and climate chamber were conducted to assess susceptibility to several SCN populations. The primary population used for both the field inoculation (Tonglu 2024) and the main greenhouse resistance assays was HG Type 0 (Race 3). Additional resistance assays were conducted in the climate chamber using Race 1 (HG 2.5.7), Race 2 (HG 1.2.5.7), and Race 5 (HG 2.5.7) (Figure 4E-G). Nematode cultures were maintained and used according to established protocols with slight modifications (Huang et al. 2021). For greenhouse assays, hatched second-stage juveniles (J2s) obtained from crushed cysts were collected for inoculation. A suspension containing 1,000 J2s in 1 mL was inoculated onto roots of 10-day-old plants at 26 °C. The use of J2s at a standardized density allows precise control of infection under uniform conditions, which is suitable for quantifying early infection events. In contrast, field and chamber screenings were conducted with cysts/eggs at higher inoculum densities to ensure robust infection establishment under variable soil and environmental conditions. The number of nematodes inoculated at different stages was counted, and fuchsin-stained roots were observed using an Olympus BX53 microscope (Olympus, Japan). Each experiment included more than four plants per treatment and was repeated at least three times.

### Genotypic and Phenotypic Analysis of *lmm3* Heterozygotes

A total of 154 seeds from a heterozygous genotype (*LMM3*^+/-^) plant were sown in the field in Changchun in 2023, yielding 154 F_2_-generation plants. Genotyping was done by PCR and Cleaved Amplified Polymorphic Sequences (CAPS) methods (Konieczny and Ausubel 1993). Thirty-nine plants were homozygous for *lmm3*, aligning with its lesion-mimic phenotype. Of the 115 plants showing a wild-type phenotype, six were disregarded due to inconclusive genotyping. CAPS digestion identified 73 plants as heterozygous and 36 as homozygous wild type.

Agronomic traits including 100-seed weight, seed-weight per plant, and plant height, were evaluated for each genotype. Plant height was determined at the R8 stage (full maturity). For each genotype, 10 plants were sampled per replication, with three replications, totaling 30 plants per genotype. Each plant was treated as a biological replicate, and mean values per genotype were used for subsequent statistical analysis.

### Bulked-Segregant Analysis (BSA) and Genetic Mapping

Bulked-segregant analysis was conducted using the BC_1_F_2_ segregating population in Changchun, China (43.88°N, 125.35°E) in 2023, which was generated by backcrossing *lmm3* with Williams 82. DNA was extracted from 39 F_2_ individuals exhibiting the *lmm3* mutant phenotype and grouped into a mutant pool. Library construction and bulked-segregant analysis followed methods described previously (Feng et al. 2019; Ye et al. 2025). Sequence data were deposited in the Genome Sequence Archive (GSA) database in the BIG Data Center (https://bigd.big.ac.cn/gsa/index.jsp) under accession number CRA012636 and have also been deposited in the NCBI Sequence Read Archive (SRA) under accession number SRR35145381 and the corresponding BioProject accession PRJNA1311168.

For map-based cloning, F_2_ plants from a cross between *lmm3* mutant and Hedou 12 were selected. Hedou 12 was chosen as the parental line to provide sufficient genetic polymorphisms for mapping. Plants exhibiting a necrotic-leaf phenotype were chosen for preliminary mapping using previously reported molecular markers between Hedou 12 and Williams 82 (Song et al. 2015). Insertion or deletion (INDEL) molecular markers were designed and developed for further fine mapping of the *LMM3* locus (Table S2).

### Soybean Stable Transformation

For mutation complementation, the wild-type *LMM3* coding sequence from Williams 82 was fused to green fluorescent protein (GFP) and placed under the control of a 3.12-kb *LMM3* promoter fragment. This construct was cloned into the pCAMBIA3301 vector between *Spe* I and *EcoR* I restriction sites, generating the *LMM3_pro_:LMM3-GFP* plasmid. The plasmid was introduced into *Agrobacterium tumefaciens* strain EHA105 and used to transform cotyledonary explants from the *lmm3* mutant soybean, following a modified transformation protocol based on Yamada et al (Yamada et al. 2010).

The resulting T_0_ transgenic seedlings were initially screened for positive transformants by appling a 1/1000 (v/v) dilution of Bar herbicide to the leaves using a cotton swab. Putative transgenic plants were then validated through PCR amplification of the *Bar* gene using specific primers, with genomic DNA extracted from the leaf tissue to confirm the presence of the transgene.

### Trypan Blue and DAB Staining

To detect spontaneous cell death and H_2_O_2_ accumulation, staining assays were performed using fresh leaves. Trypan blue staining was used to detect spontaneous cell death, while 3,3’-diaminobenzidine (DAB) staining was employed to assess H_2_O_2_ accumulation following previously described procedures (Wang et al. 2020).

### Vector Construction and Generation of Transgenic Hairy Roots

The SCN-responsive *HIP* promoter (1 kb) was cloned and fused with the *SNAP18_lmm3_* coding sequence using the Golden Gate Modular Cloning (MoClo) toolkit. Briefly, the *HIP* promoter, the SNAP18_lmm3_ ORF, and the nopaline synthase (NOS) terminator were assembled into a Level 1 transcriptional unit. This unit was then integrated into the Level 2 binary vector pAGM4673 alongside a selection cassette. The selection cassette consisted of a green fluorescent protein (GFP) reporter gene driven by the double CaMV 35S promoter with the TMV omega enhancer (pICH51288). All assembly steps were performed via Golden Gate cloning as previously described (Weber et al. 2011).

The resulting binary construct, pAGM4673-*HIP*::*SNAP18_lmm3_*, was transformed into *Agrobacterium rhizogenes* strain ArQua1. Transgenic soybean hairy roots were generated according to the protocol described by Chen et al (Chen et al. 2024). After 2∼3 weeks of induction, emerging hairy roots were screened for successful transformation by visualizing GFP fluorescence using a stereomicroscope. Only robustly fluorescing root lines were selected for subsequent SCN infection assays and molecular characterization.

### Statistical Analysis

All experiments were conducted with at least two independent biological replicates. Error bars in the figures represent either the standard deviation (SD) or the standard error of the mean (SEM), as indicated in each figure legend. Statistical analyses were performed using GraphPad Prism software (version 10.2.0).

To assess statistical significance, appropriate tests were selected based on data distribution and experimental design. These included two-tailed Student’s *t*-tests for normally distributed data, and one-way or two-way ANOVA for multi-group analyses. A *P*-value < 0.05 was considered statistically significant. Statistical significance is denoted as follows: Asterisks indicate differences from two-tailed Student’s *t*-tests (\**P* < 0.05; \*\**P* < 0.01; \*\*\**P* < 0.001). Distinct letters represent significant differences for one-way ANOVA or two-way ANOVA analyses.

## Acknowledgements

We thank Dr. Shiming Liu from Chinese Academy of Agricultural Sciences for providing Forrest seeds. This work was supported by the National Natural Science Foundation of China (32488102 to X.F., 32301809 to D.W., and 32272478, 32102146 to S.H.), National Key Research and Development Program of China (2023YFD1401000, 2023YFD1400400 to S.H.), Science and Technology Development Plan Project of Jilin Province of China (20240602056RC to D.W.), Innovation Team Project of Northeast Institute of Geography and Agroecology, Chinese Academy of Sciences (2022CXTD03 to D.W.).

## Author Contributions

X.F. conceived and directed the project. S.Y. guided the details of the project and revised the manuscript. D.W. and S.H. designed and performed most of the experiments. D.W., S.H., S.Y. and X.F. drafted the manuscript with approval of the version to be submitted from all authors. L.W., L.W. and H.Y. performed wet-lab experiments. L.C., and M.H. conducted nematode-inoculation experiments. C.L. provided SCNs. All authors read and approved of its content.

## Conflicts of Interest

The authors declare no competing interests.

## Supporting Information

**FIGURE S1.**
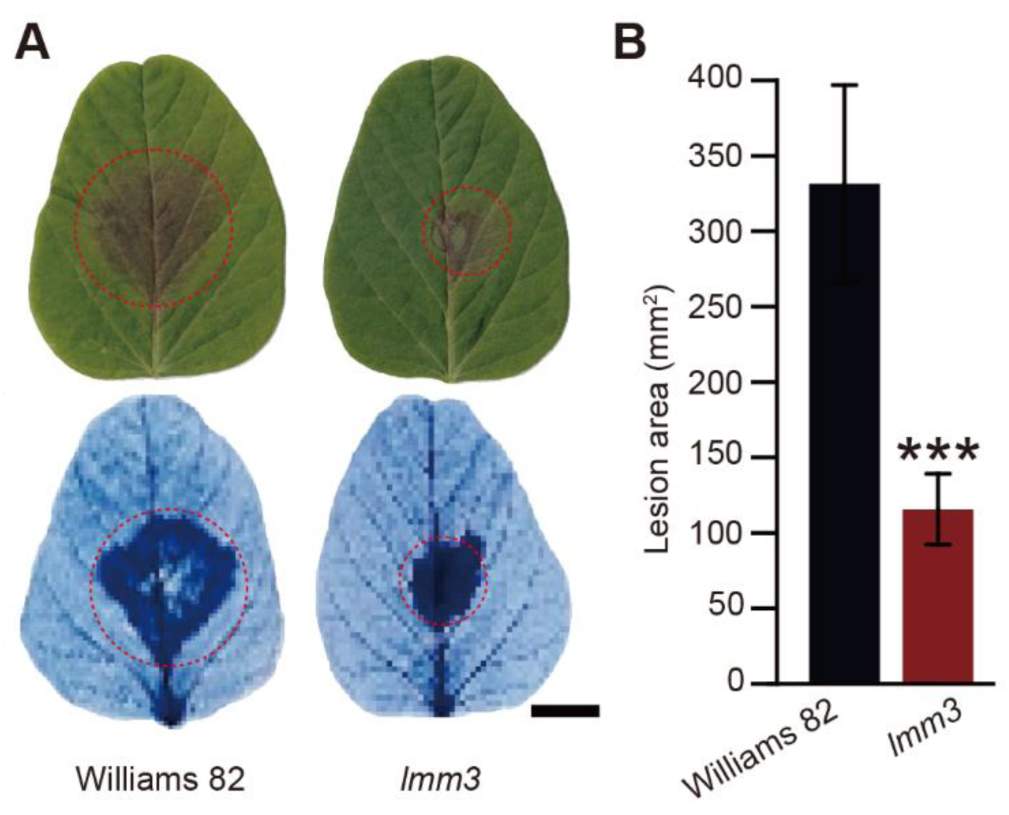
The *lmm3* mutant has reduced susceptibility to *Phytophthora sojae* stain P7076. (A) Infection phenotypes of Williams 82 and the *lmm3* mutant at 48 hours post-inoculation (hpi) with *Phytophthora sojae* strain P7076. Trypan blue staining (circled in red) revealed that infection levels were similar to those of the leaves shown above. Scale bar = 1 cm. (B) Quantification of lesion areas on leaves as in (A). Data are the mean ± SD of six biological replicates. Asterisks indicate statistically significant differences between means (*** *P* < 0.001, Student’s *t*-test).

**FIGURE S2.**
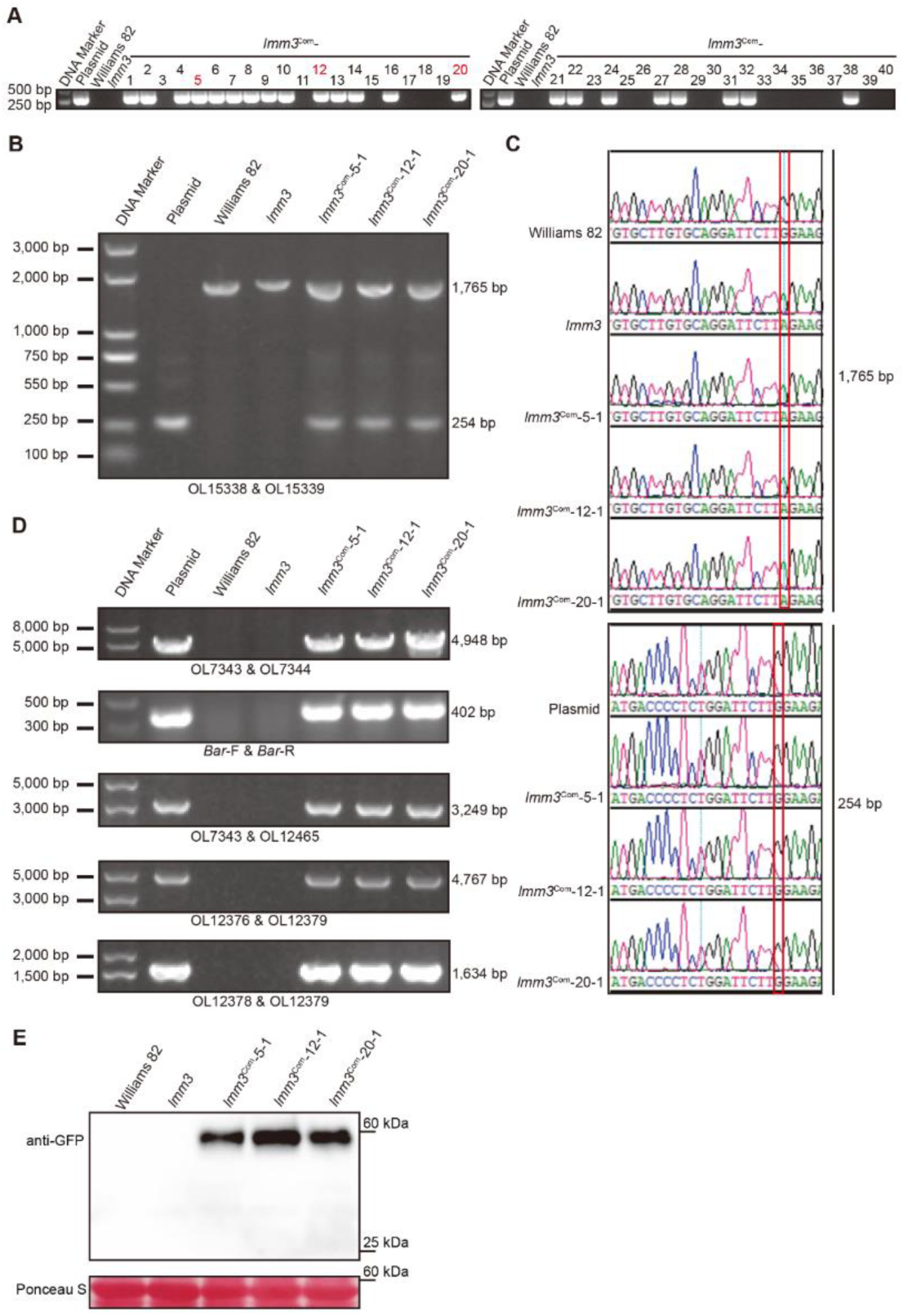
Genotypic analysis of transgenic *lmm3 LMM3_pro_:LMM3–GFP* complementation lines. (A) PCR amplification of the *Bar* gene to detect T_0_-generation transgenic *lmm3* (*LMM3_pro_:LMM3*–*GFP*) complementation lines. Forty plants were tested, designated as *lmm3*^Com^-1 to *lmm3*^Com^-40. (B) PCR verification Williams 82, *lmm3*, and the three independent T_1_ generation transgenic complementation plants using the primers OL15338 and OL15339. A 254-bp band was generated by the *Glyma.18G022500* coding sequence in a plasmid, a 1,765-bp band was produced by amplification of genomic DNA in Williams 82 and *lmm3*, and both a 254-bp band and a 1,765-bp band were amplified in the three transgenic complementation lines. (C) DNA sequencing to verify the genotype of plants analyzed in (B). The upper and lower panels show the *Glyma.18G022500* sequence, indicating the transgenic events. Red boxes represent the mutation site of the *lmm3* mutant in the gDNA (upper panel) and the corresponding successfully complemented mutation in the plasmid used for transformation and the corresponding recovered transgenic plants (lower panel). (D) PCR genotyping to identify transgenic complementation events. PCR was carried out using the primers indicated to amplify the T-DNA region of the complementation vector, wild-type Williams 82, *lmm3* and the three independent transgenic complementation lines. (E) Western blot with anti-GFP antibodies to detect LMM3*–*GFP accumulation in transgenic *lmm3* complementation lines. Ponceau S was used as a loading control.

**FIGURE S3.**
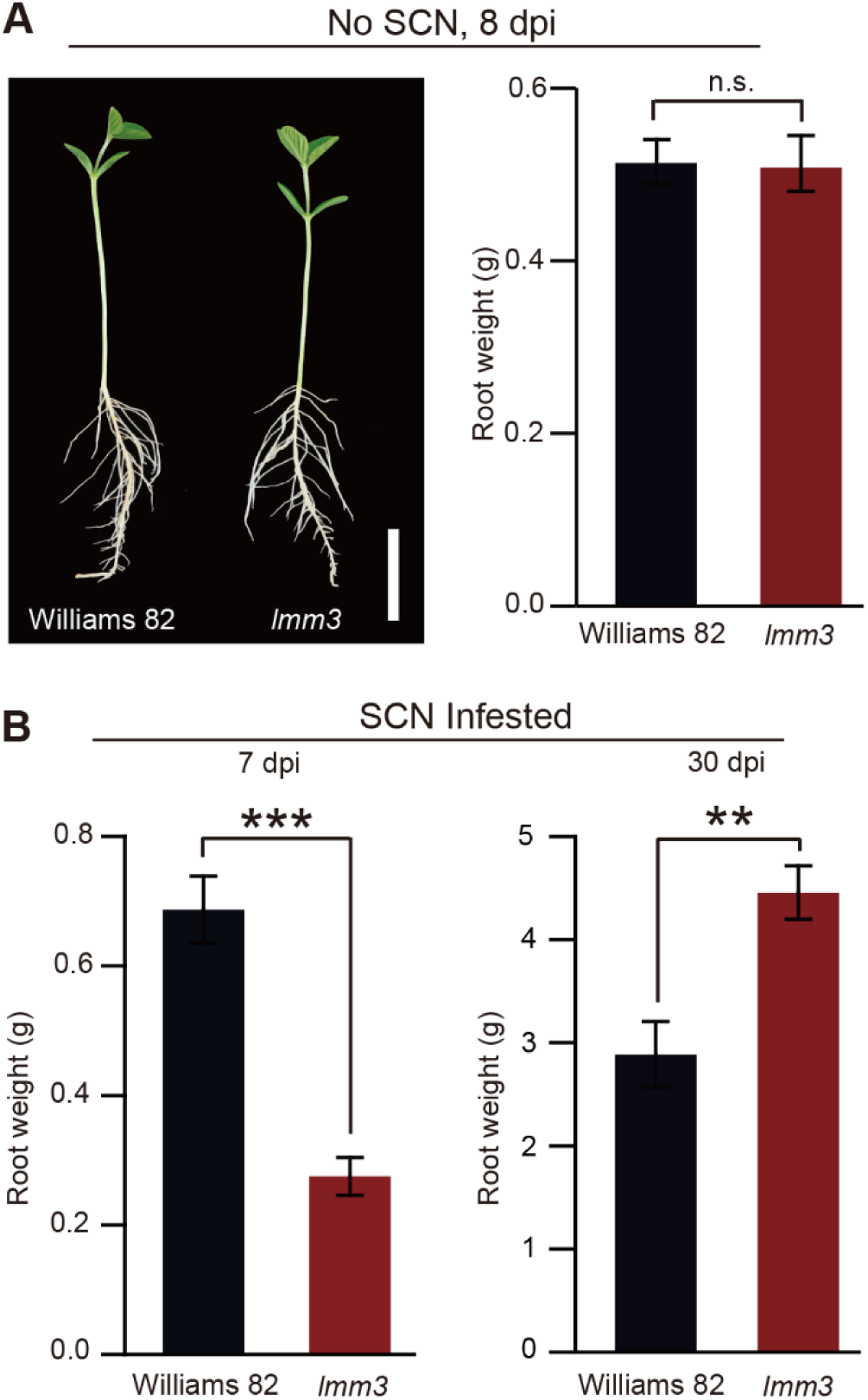
Analysis of Williams 82 and *lmm3* root phenotypes. (A) Left: Williams 82 and *lmm3* mutant at 8 DAG grown in vermiculite. Scale bar = 5 cm. Right: Root fresh weight of Williams 82 and *lmm3* at 8 DAG grown in vermiculite. Data are the mean ±SD of n = 10 plants. n.s. indicates a statistically non-significant difference (Student’s *t*-test). (B) Root fresh weight for Williams 82 and *lmm3* at 7 and 30 dpi with SCN. Data are the mean ±SEM of n = 10 plants. ** *P* < 0.01, *** *P* < 0.001 (Student’s *t*-test). All the experiments were repeated three times with consistent results.

**FIGURE S4.**
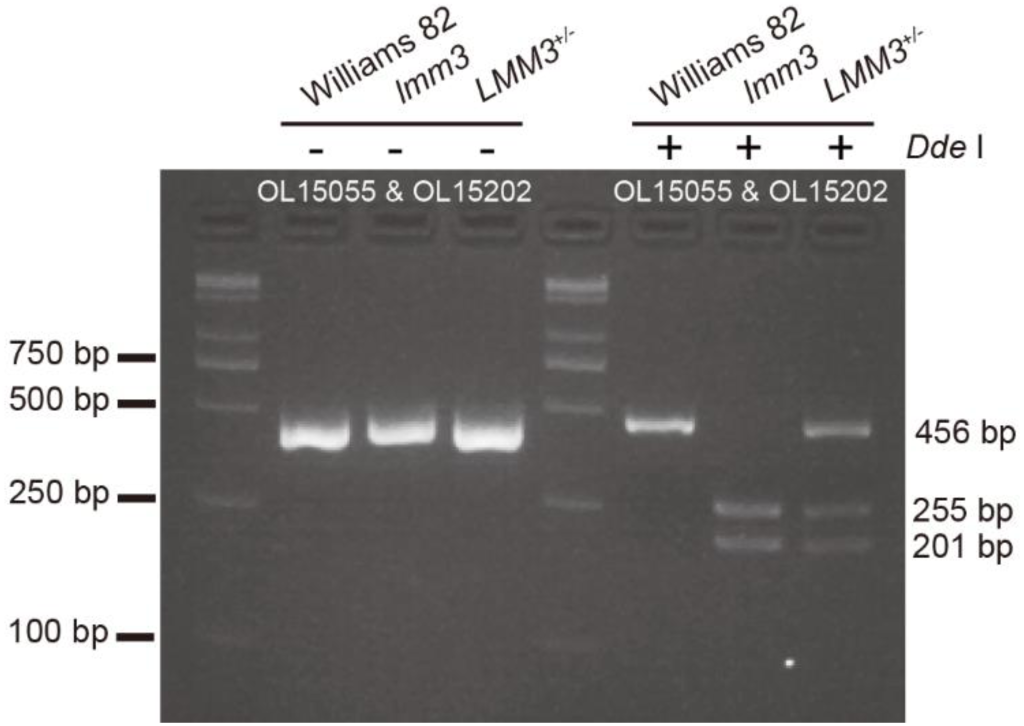
CAPS genotyping of Williams 82, the *lmm3* mutant and heterozygous *LMM3^+/-^*plants. Primers spanning the *lmm3* mutation site were used, followed by *Dde*I endonuclease digestion to distinguish genotypes.

**FIGURE S5.**
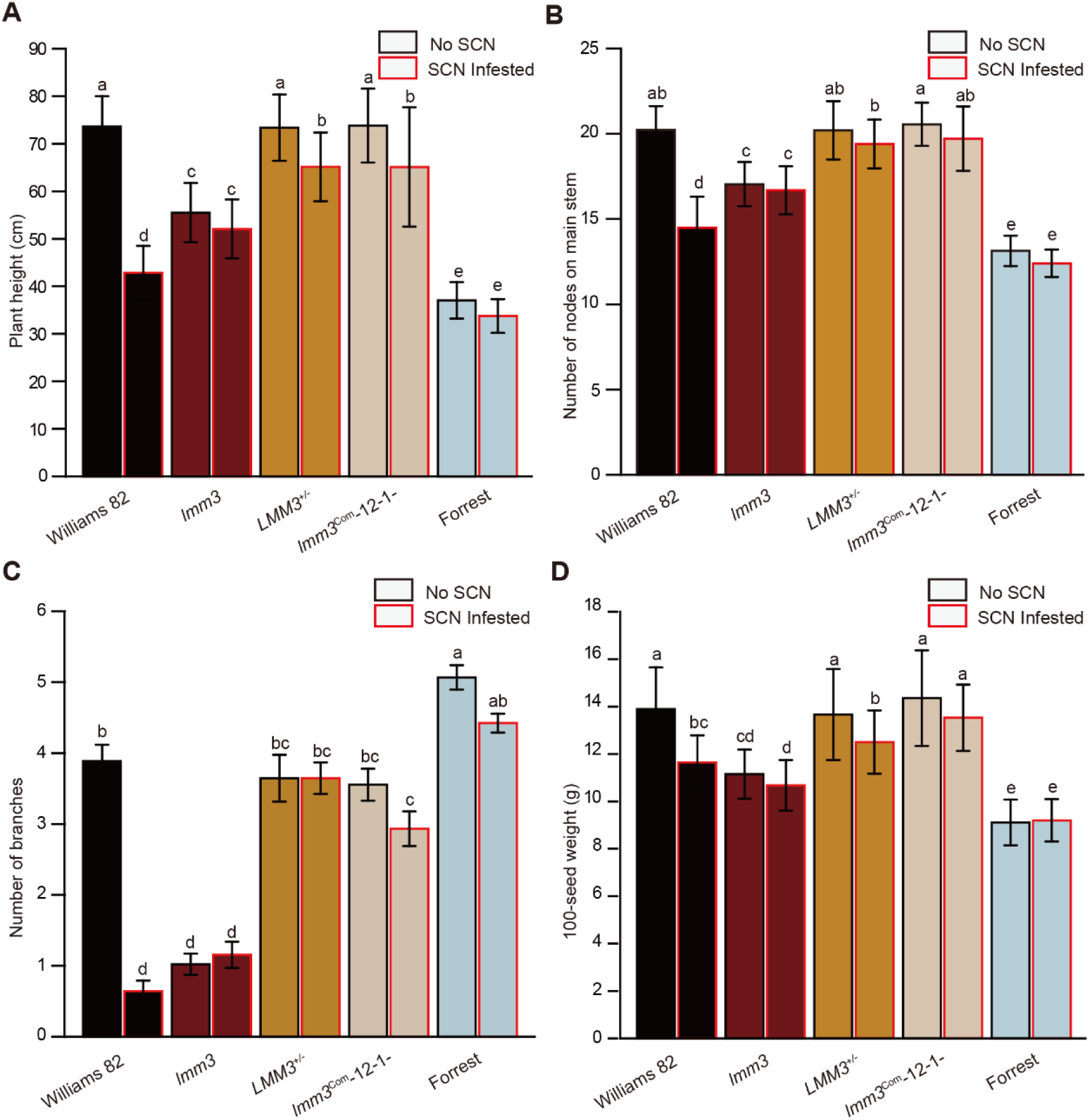
Yield metrics for field-grown Williams 82, *lmm3* mutants, *LMM3^+/-^* heterozygous plants, T_2_ transgenic complementation plants (*lmm3*^Com^-12-1-, carrying *LMM3_pro_:LMM3–GFP* constructs in the *lmm3* mutant background) and Forrest, with or without SCN infestation. (A–D) Plant height (A), number of nodes on main stem (B), number of branches (C) and 100-seed weight (D) for data collected in Tonglu in 2024. Data are presented as mean ±SD for panels A, B and D, and as mean ±SEM for panel C (n = 45 plants) Statistical significance was determined using two-way ANOVA, with different letters indicating *P* < 0.05.

**TABLE S1.** Yield comparison (corrected to 13% moisture) for five genotypes under SCN-inoculated and non-inoculated conditions in Tonglu, 2024.

|  |  | No SCN |  |  |  |  | SCN infested |  |  |  |  |
| --- | --- | --- | --- | --- | --- | --- | --- | --- | --- | --- | --- |
| Variety |  | Williams | *Imm3* | *LMM* | *Imm3*<sup>Com</sup> | Forrest | Williams | *Imm3* | *LMM3* | *Imm3*<sup>Com</sup> | Forrest |
|  |  | 82 |  | 3<sup>+/-</sup> | -12-1- |  | 82 |  | +/- | -12-1- |  |
| Seed yield (kg) | Plot 1 (12 rows) | 6.99 | 3.10 | 6.78 | 6.92 | 3.14 | 1.51 | 2.46 | 5.48 | 5.52 | 2.90 |
|  | Plot 2 (12 rows) | 6.69 | 3.05 | 7.16 | 7.03 | 3.26 | 1.47 | 2.40 | 5.61 | 5.57 | 2.75 |
|  | Plot 3 (12 rows) | 7.05 | 3.23 | 6.83 | 6.83 | 3.35 | 1.48 | 2.53 | 5.56 | 5.69 | 2.80 |
| Seed yield per plot (kg) |  | 6.91 | 3.13 | 6.92 | 6.93 | 3.25 | 1.49 | 2.46 | 5.55 | 5.59 | 2.82 |
| Seed yield (kg/ha) with ~87,500 plants/ha |  | 2399.31 | 1086.81 | 2402.78 | 2406.25 | 1128.47 | 517.36 | 854.17 | 1927.08 | 1940.97 | 979.17 |

Note: All yield data were corrected to a standard moisture content of 13% using a KETT (Japan)
PM8188A Moisture Meter. Plots were harvested by hand, with edge rows excluded to minimize
border effects; the remaining 12 rows per plot were threshed manually on the same day. The
experiment was conducted using a randomized complete block design with three blocks and equal
replication per genotype under both SCN-inoculated and non-inoculated conditions.

**TABLE S2.**
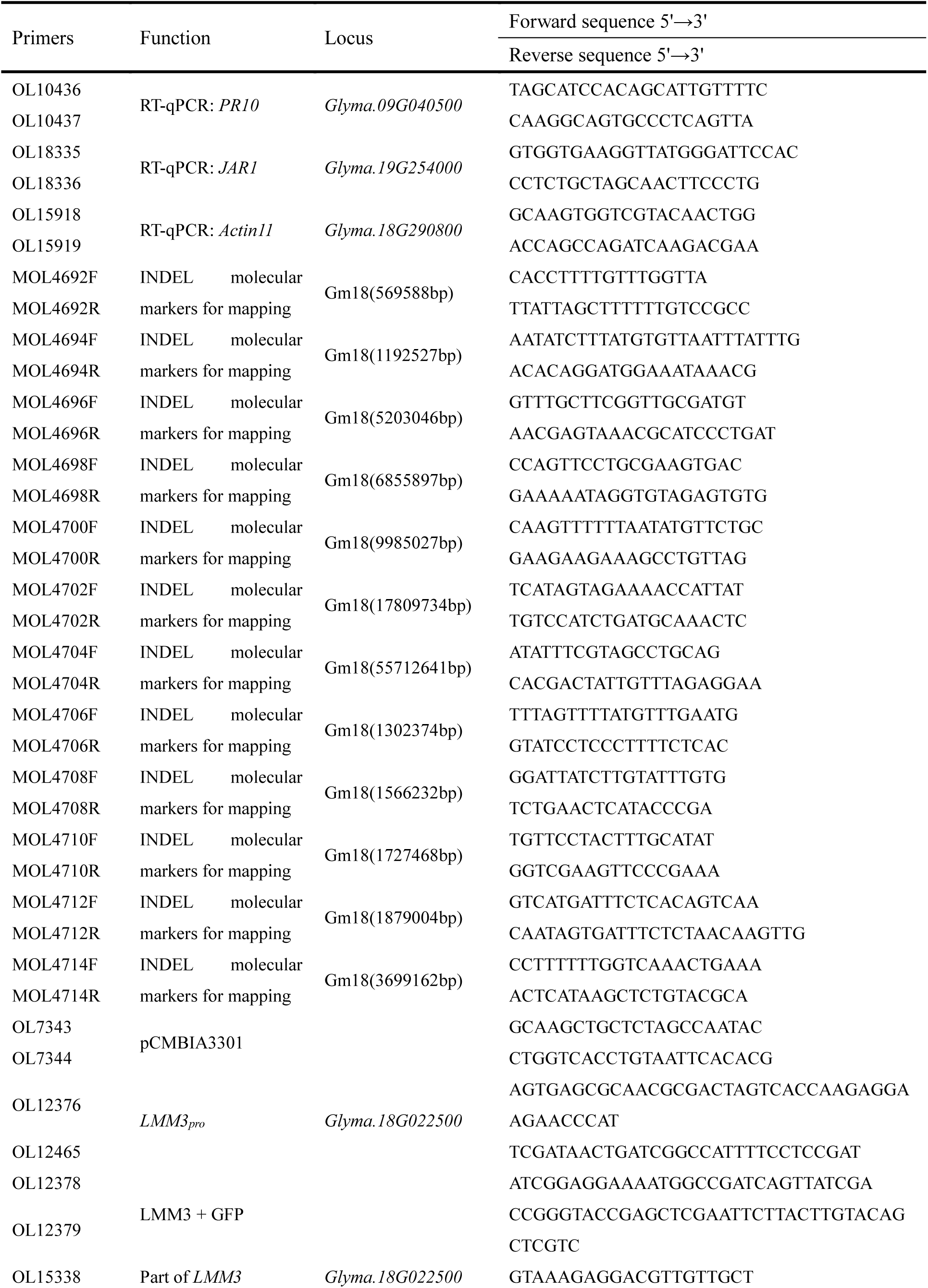
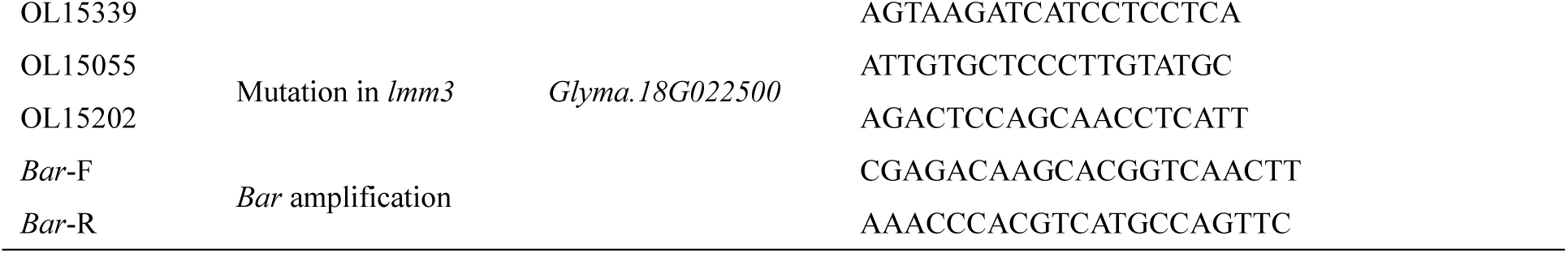
Primers used in this study.

## Notes

https://bigd.big.ac.cn/gsa/index.jsp

https://www.ncbi.nlm.nih.gov/sra/

https://www.ncbi.nlm.nih.gov/bioproject/

### Competing Interest Statement

The authors have declared no competing interest.

